# Biomechanical dependence of SARS-CoV-2 infections

**DOI:** 10.1101/2022.02.17.479764

**Authors:** Alexandra Paul, Sachin Kumar, Tamer S. Kaoud, Madison Pickett, Amanda L. Bohanon, Janet Zoldan, Kevin N. Dalby, Sapun H. Parekh

## Abstract

Older people have been disproportionately vulnerable to the current SARS-CoV-2 pandemic, with an increased risk of severe complications and death compared to other age groups. A mix of underlying factors has been speculated to give rise to this differential infection outcome, including changes in lung physiology, weakened immunity, and severe immune response. Our study focuses on the impact of biomechanical changes in lungs that occur as individuals age, *i.e.*, the stiffening of the lung parenchyma and increased matrix fiber density. We used hydrogels with an elastic modulus of 0.2 and 50 kPa and conventional tissue culture surfaces to investigate how infection rate changes with parenchymal tissue stiffness in lung epithelial cells challenged with SARS-CoV-2 Spike (S) protein pseudotyped lentiviruses. Further, we employed electrospun fiber matrices to isolate the effect of matrix density. Given the recent data highlighting the importance of alternative virulent strains, we included both the native strain identified in early 2020 and an early S protein variant (D614G) that was shown to increase the viral infectivity markedly. Our results show that cells on softer and sparser scaffolds, closer resembling younger lungs, exhibit higher infection rates by the WT and D614G variant. This suggests that natural changes in lung biomechanics do not increase the propensity for SARS-CoV-2 infection and that other factors, such as a weaker immune system, may contribute to increased disease burden in the elderly.

## Introduction

The novel coronavirus disease first reported in 2019 (COVID-19) attained global pandemic status in mid-April 2020 and is associated with severe acute respiratory distress symptoms [1]. Global statistics on SARS-CoV-2 infection have shown that it causes severe lung damage, particularly among older adults compared to other segments of the population, causing 23-fold increased mortality in more senior members of society than younger individuals [2]–[4]. The age-dependent pattern of SARS-CoV-2 infection severity, especially in older people with lung comorbidities, has emerged as a significant global risk factor [5]. Despite the statistically significant increased disease burden in older people, the reasons for age-dependent SARS-CoV-2 infection severity are unclear. Hypotheses have suggested that a cytokine “storm”, hyperactivity of the immune system upon detecting the infection, or an inherently weaker immune system may lie at the heart of the increased burden in older people [6]–[8]. While the host immune response certainly plays an important role, it is also necessary to study underlying factors that influence the initial virus infection path to map risk factors and inform treatment options. The current study focuses on quantifying how the underlying biomechanical differences in lung parenchyma of the elderly and young affect the propensity of SARS-CoV-2 infection.

Physiological studies of lung tissue from subjects of different ages have shown that lung properties change during aging, e.g., more fibrotic tissue is present in lungs of older patients, and lung parenchymal tissue compliance decreases with age in addition to several biochemical changes [9]– [11]. As the parenchymal lung tissue stiffens, on account of changes in the extracellular matrix (ECM) composition and architecture, it can have substantial effects on cellular function, similar to what is seen in other tissues [12], [13]. ECM fiber density and composition have been shown to significantly increase with lung pathology at any life stage, as seen with asthma or idiopathic pulmonary fibrosis, among others [14]. Indeed, more fibrotic and stiffer lungs are more susceptible to environmental exposures and infectious pathogens, such as viruses [15]. Therefore, lung parenchymal tissue stiffness is a variable that could influence viral infection by SARS-CoV-2 and is a relevant physiological difference in elderly versus younger segments of the population that might partially explain differences in infection severity with age.

ECM stiffness is now well known to regulate cell physiology and behavior, causing cells to reorganize and remodel their membrane and cytoskeletal structures [16]–[18]. For example, membrane permeability, which plays a critical role in cellular uptake of biomolecules, is altered by changes in substrate stiffness [19], [20]. A few studies have shown how substrate stiffness regulates the cellular uptake of micelles and nanoparticles in cancer and disease models [21], [22]. Moreover, studies with adenoviruses and influenza viruses have shown how initial viral entry is regulated by mechanical cues on the cell surface, further suggesting a prominent role of viral and tissue mechanics in the infection process [23]–[27]. Finally, ECM stiffness and microenvironment properties have been found to affect the ability of stromal cells to induce chemokines and cytokines in the recruitment and differentiation of monocytes to control the innate immune system against lung diseases [28], [29]. However, the impact of biomechanical changes in the local tissue environment on the infection of cells by viruses remains largely unexplored.

Viral internalization depends on both the characteristics of the virus and the host cell properties (which are known to change with local tissue properties). Model-based studies have shown that early-stage invagination of three different viruses (reo-, adeno-, papilloma-, and polio-viruses) depends on contact pressure and force exerted on the host cell membranes [23]. Initial contact pressures depended on the virus geometry and virus and cell stiffness [30]. Although several coronaviruses and influenza viruses have been known to disproportionally affect older people, the precise reasons are still unknown [3], [31]. Thus, understanding and, eventually, controlling the early viral entry in the lungs could be one of the best tactics to manage SARS-CoV-2 infection in older patients.

In the present study, we experimentally mimic age-related variation in lung stiffness by culturing Calu-3 airway epithelial cells on surfaces with different elastic moduli (0.2 kPa, 50kPa, 100000 kPa). A similar range (0.2 – 50 kPa) has been found in healthy *versus* fibrotic tissue in lungs derived from patients [9] and has been used to model these conditions *in vivo* [32]–[34]. Using pseudotyped lentiviruses (PV) expressing the SARS-CoV-2 Spike (S) protein, we evaluated how the biomechanical compliance of the tissue affects SARS-CoV-2 infection in lung airway epithelial cells. We further studied infection dynamics on electrospun fibers that were sparse or dense, mimicking ECM densification in the aging lung.

## Materials and Methods

### Plasmids

Plasmids used to express components of the HIV virion under the CMV promoter (HDM-Hgpm2, pRC-CMV-Rev1b, and HDM-tat1b) were obtained from BEI resources as NR-52516, NR-52519, and NR-5251, respectively [35]. Plasmids for lentiviral backbone expressing fluorescent reporter under CMV promoter (pHAGE2-CMV-ZsGreen-W) or human ACE-2 gene (GenBank ID NM_021804) under an EF1a promoter (pHAGE2-EF1aInt-ACE-2-WT) were obtained from BEI resources as NR-52520 and NR52512, respectively [35]. The envelope vector expressing VSV-G (vesicular stomatitis virus glycoprotein) was obtained from Cell Biolabs (pCMV-VSV-G, Part No. RV-110). To generate the expression plasmid for D614G mutant of SARS-CoV-2 Spike (S) protein, a plasmid expressing a codon-optimized WT S (Genbank ID NC_045512) under a CMV promoter was obtained from BEI Resources (HDM-IDTSpike-fixK, NR-52514) [35] and employed as a template for site-directed mutagenesis.

### Generation of HEK293T cells with stable expression of human ACE-2

HEK293T cells were transduced with a lentiviral vector (pHAGE2-EF1aInt-ACE-2-WT) that expresses human ACE-2 under an EF1a promoter. A clone that expresses a high level of ACE-2 was selected based on its ability to be highly infected by the pseudotyped lentiviral (PVs) particles bearing WT-SARS-CoV-2 S protein (ZsGreen reporter) and was confirmed using western blotting.

### Generation of (SARS-CoV-2) S protein Pseudotyped Lentivirus

PVs were prepared by co-transfecting 293T cells with plasmids for (1) lentiviral backbone containing fluorescent reporter (pHAGE2-CMV-ZsGreen-W), (2) HIV virion formation proteins under CMV promoters (HDM-Hgpm2, pRC-CMV-Rev1b, and HDM-tat1b; and (3) viral entry proteins SARS-CoV-2 S protein (WT, D614G mutant) or VSV G as a positive control for infectivity. A detailed protocol for creating these PVs was reported by Crawford *et al.* [35]. The lentiviral particles were harvested by collecting and filtering the HEK 293T cell supernatant at 60-72 h post-transfection. The collected virus was frozen at −80°C as fractions. Successful viral entry was confirmed and quantified by measuring the ZsGreen expression signal in infected cells. Viral titers were determined following a previously published protocol with few modifications [35]. In summary, 12500 ACE-2 HEK 293T cells were seeded per well of 96 well plates (39000 cells per cm^2^, black with clear bottom, coated with Poly-L-lysine D). Cells were allowed to adhere 24 hours before treatment with a serial dilution of the lentiviral particles in full media containing 5 μM polybrene (in triplicate). 60-72 hours post-infection, cells expressing ZsGreen, and the total number per well were estimated using the lncuCyte® ZOOM equipment with a ×10 objective. Viral titers were calculated using the Poisson formula.

### Electrospun fiber mat fabrication and hydrogels

Polycaprolactone (PCL) fibers were prepared using a horizontal electrospinning set-up. A 10 wt% solution of PCL in Hexafluoro-2-propanol (HFIP) was prepared and loaded into a 10 mL syringe with a 25G needle. The tip to collector distance was set to 11 cm, the flow rate 1.0 ml/h, with an applied voltage of 10 kV. Coverslips of 12 mm diameter were plasma cleaned for one minute at 25mA using the EMITECH K100X and subsequently attached to the rotating collector using double-sided tape. Both microenvironments were fabricated at a collector speed of 310 rpm. The duration of fabrication for the sparse and dense microenvironment was seven minutes and fifteen minutes, respectively. In preparation for cell culture, the scaffolds were coated with 50 μg/ml collagen I in a complex Calu-3 medium (see below) at 37°C overnight.

Commercial hydrogels coated with collagen I monomers with two different stiffnesses were used for hydrogels: 0.2 kPa and 50 kPa (Softview™, Matrigen). The hydrogels were used without further modification.

### Scanning electron microscopy

The sparse and dense electrospun scaffolds were mounted on SEM stubs using double-sided conductive tape. The samples were coated with 7.0 mm Pt/Pd using a Cressington 208Hr Sputter Coater. After sputtering, the samples were loaded into the Zeiss Supra 40VP SEM for subsequent imaging. Images were acquired under vacuum and at magnifications ranging from 0.5 to 1.5 kX.

### Cell culture and viral infection

Experiments were performed using lung adenocarcinoma cells (Calu-3, ATCC HTB-55™) obtained from the American Type Culture Collection (ATCC) and on human embryonic kidney cells (HEK293T, ATCC® CRL-3216™) overexpressing ACE-2. Calu-3 cells were maintained in EMEM (with EBSS and L-glutamine, BioWhittaker, Lonza) supplemented with 20% FBS (heat-inactivated, Gibco), 0.1 mM MEM non-essential amino acids (Gibco), and 10 mM HEPES (Gibco) at pH 7.4. The passage number was kept below 10 for all experiments, and cells were incubated at 37°C in a 5% CO_2_ atmosphere. ACE-2 HEK 293T cells were cultured in DMEM + GlutaMax™-I medium (Gibco) supplemented with 10% FBS. The passage number was below 25 for all experiments.

Cells were either seeded on tissue culture polystyrene surfaces (TCPS) at 125,000 cells per cm^2^ or on hydrogels or fibers at 187,500 cells per cm^2^. Cells were allowed to attach for 2 days. Then all samples were exposed to the same volume of PVs (1:4 dilution from stock, 16.6 μl/10,000 cells) encoding for ZsGreen transcription in a complex medium containing 5μg/ml polybrene (Merck Millipore). Using qPCR, we confirmed the exposure to be 8.77E6 IU/ml for WT and 2.25E6 IU/ml for D614G (**Supplementary Figure S1 and Table S1**). The infection/expression period was allowed to progress for 4 days, after which cells were washed with warmed PBS, stained with 5 μM DRAQ5 (Thermo Scientific, nuclear dye, 5 minutes at 37°C), fixed in 4% paraformaldehyde for 20 minutes, and washed once more in PBS. Samples were stored in PBS at 4°C until analysis.

### ACE-2 receptor expression

Expression of ACE-2 in Calu-3, on TCPS and hydrogels, and ACE-2 HEK 293T on TCPS was evaluated using immunofluorescence as shown previously [36]. Cells were seeded, left to adhere for 48 hours, fixed as described above, blocked with 5% BSA for 1h at room temperature, repeatedly rinsed with PBS, and incubated with 4 μg/ml Anit-ACE-2 conjugated to Alexa 647 (Santa Cruz Biotechnology, SC-390851) at 4°C overnight. The next day, the sample was washed several times with PBS and stored in PBS at 4°C until analysis.

### Confocal microscopy acquisition and data analysis

An inverted confocal microscope (Olympus, FV3000) was used to quantify ZsGreen, DRAQ5, and Anti-ACE-2-Alexa647 fluorescence. Images were acquired using a 20x air immersion objective with 0.45 NA (for cells on TCPS) or 0.75 NA (for cells on hydrogels) (both Olympus). ZsGreen was excited with a 488 nm laser line and detected between 500-540 nm, and DRAQ5 was excited at 640 nm and detected between 650-750 nm. Anti-ACE-2-Alexa647 fluorescence in separate samples was excited at 640 nm and detected between 650-750 nm. The confocal aperture was kept at 1 Airy Disk for all experiments.

The ZsGreen expression area was quantified and normalized to the cell area estimated from the DRAQ5 staining using a custom ImageJ script. To estimate the cell area, a Gaussian blur filter with sigma 10 pixels was applied to a max intensity projection of the DRAQ 5 channel, followed by applying the “default” ImageJ threshold. The ZsGreen channel was treated similarly except that the blur sigma was 2 pixels, and the threshold was set at 98.5% of the channel’s mean fluorescence intensity. A manual ROI selection was performed from the brightfield channel followed by a Gaussian blur with sigma 0.5 pixels for the antibody expression. Then the mean fluorescence intensity per ROI was measured. To account for the variance in viruses/mL, the numbers for D614G were multiplied by 3.89 (D614G titer/WT titer). To further account for the experimental difference, we only compare WT and D614G on TCPS to verify our results against existing literature.

### Statistical analysis

Statistical differences were analyzed using GraphPad Prism. Outliers were removed using ROUT with Q=1%. For experiments containing 2 groups, an unpaired two-sided t-test was used to determine statistical significance (p < 0.05). For experiments with 3 or more groups, an ordinary one-way ANOVA was followed by Tukey’s multiple comparison test. N denotes the number of biological replicates, which refers to repeated cell seedings on different days. n denotes the number of images analyzed.

## Results and Discussion

### SARS-CoV-2 pseudotyped lentiviruses infect Calu-3 lung cells

Upon entering a host cell, viruses rely on the host cell machinery and metabolism to replicate their genetic material and reproduce themselves. To initiate this process, SARS-CoV-2 infects host cells by binding to the Angiotensin-converting enzyme (ACE-2) receptor using its receptor-binding domain of the Spike (S) protein that is displayed as part of the protein corona on the virus surface [37], [38]. The ACE-2 receptor is found in epithelia linings in the body, including the lungs, intestines, and brain [39]. To explore the SARS-CoV-2 infection pathway while reducing the overall experimental risk, viral mimic particles using PVs bearing the S protein on their surfaces have been developed by numerous labs and are now available commercially. Additionally, this bioengineered system allows for the targeted introduction of key mutations. This study employs a PV system developed and published by the Bloom lab at the University of Washington, which contains a green fluorescent protein (ZsGreen) reporter in the packaged RNA to indicate successful infection. We note that this system was previously shown to have excellent specificity for infecting cells containing the ACE-2 receptor [35]. We also performed similar quality control experiments on our PV productions (**Supplementary Videos S1-S4**). This system provides a direct and quantifiable read-out of successful infectious events *via* green fluorescence, assuming that cells on all substrates translate the mRNA with similar rates.

As lung cells express a relatively high amount of ACE-2, SARS-CoV-2 causes severe lung infection and symptoms. Therefore, we used the Calu-3 lung epithelial cell line, which naturally expresses ACE-2, as a model system to study infection as a function of tissue biomechanics while at the same time evaluating infection propensity between the wild type (WT) and point mutant D614G SARS-CoV-2 variant. This allows us to determine if the infection propensity of different strains shows similar sensitivity to lung biomechanics.

Calu-3 cells were readily infected by PVs WT SARS-CoV-2 S protein (**Figure 1A, left**). Control HEK 293T cells engineered to express ACE-2 showed a similar infection pattern (**Figure 1A, Supplementary Video S2**). In contrast, no infection events were observed when the ACE-2 receptor was absent in cells (**Supplementary Video S4**). Similar to WT, we found the D614G variant infected both cell lines (**Figure 1A, right**). Quantifying the proportion of ZsGreen fluorescence relative to the total cellular fluorescence and normalizing by the viral titer for each strain, we found that D614G variants infected cells 4-fold and 3.2-fold for Calu-3 and ACE-2 HEK 293T (**Figure 1B**), respectively. These numbers align with similar measurements reporting a 2- to 10-fold on Calu-3 and ACE-2 expressing HEK 293T cells [40]–[42].

**Figure 1.**
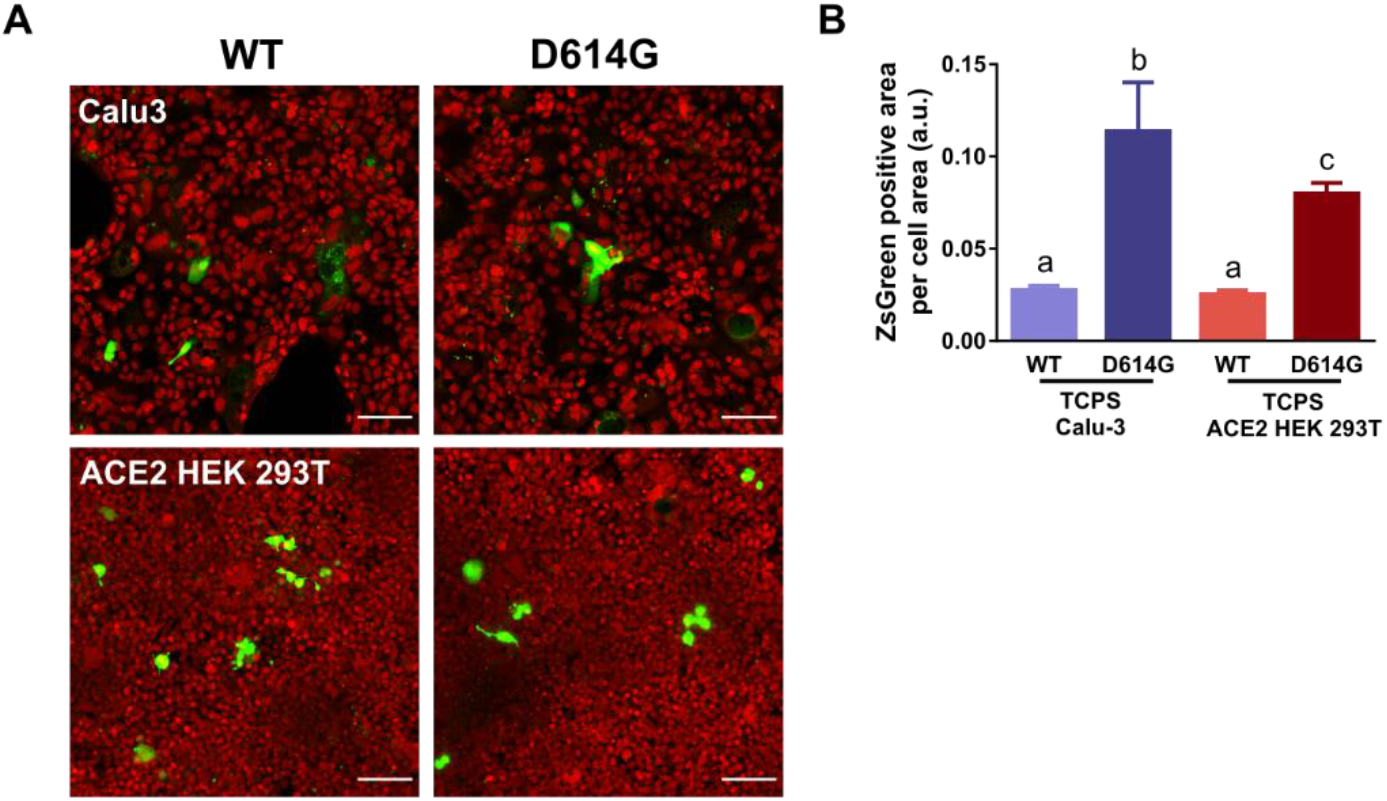
PV infection (WT and D614G) of Calu-3 lung cells and ACE-2 HEK 293T. (A) Fluorescent images showing cells (red, DRAQ5) expressing ZsGreen (green) after successful infection. Scale bars = 100 μm. (B) Increased infection rates are found with the D614G variant in both cell lines. Different letters above the bar (a-c) denote statistically significant difference (p<0.05) (mean ± SEM, N=4/n=51 for Calu3/WT, N=2/n=20 for Calu3/D614G, N=4/n=60 for ACE-2 HEK 293T/WT, N=2/n=35 for ACE-2 HEK 293T/D614G). We note that the viral titers used for the two strains were different; however, the quantification in B is normalized for the actual titer.

### PV infections are more likely on soft substrates

Since matrix stiffness affects cellular endocytosis [43], [44], tissue culture polystyrene surface (TCPS) is not an ideal experimental surface due to its unnatural stiffness of ~100000 kPa [45]. Instead, we used commercial, collagen-coated polyacrylamide hydrogels with defined elastic moduli (0.2 kPa and 50 kPa) mimicking stiffness variations found in patients [9] as substrates for Calu-3 cells to determine if infection propensity varied with tissue stiffness. Cells grown on these surfaces displayed a markedly changed morphology (**Figure 2A**) and tended to grow in smaller clusters. Using a similar quantification as in Figure 1B, both the WT and the D614G variant showed increased initial infection on *softer* surfaces (**Figure 2B**), as shown for receptor-independent uptake of metallic nanoparticles in lung cells [46]. The D614G variant infection was almost 10-fold higher than the WT PV on both substrates in these experiments. The variant was much more potent on both hydrogels than the rigid TCPS, suggesting that tuning compliance of the substrate can drastically tune the infection potential of the D614G PV. It has also been found that the SARS-CoV-2 virus is sensitive to cell stiffness using different macrophage phenotypes [47], where the virus shows higher infection rates in softer cells. In this study, we have not measured the cell stiffness, but it has been shown to match the material they are grown on [48].

**Figure 2.**
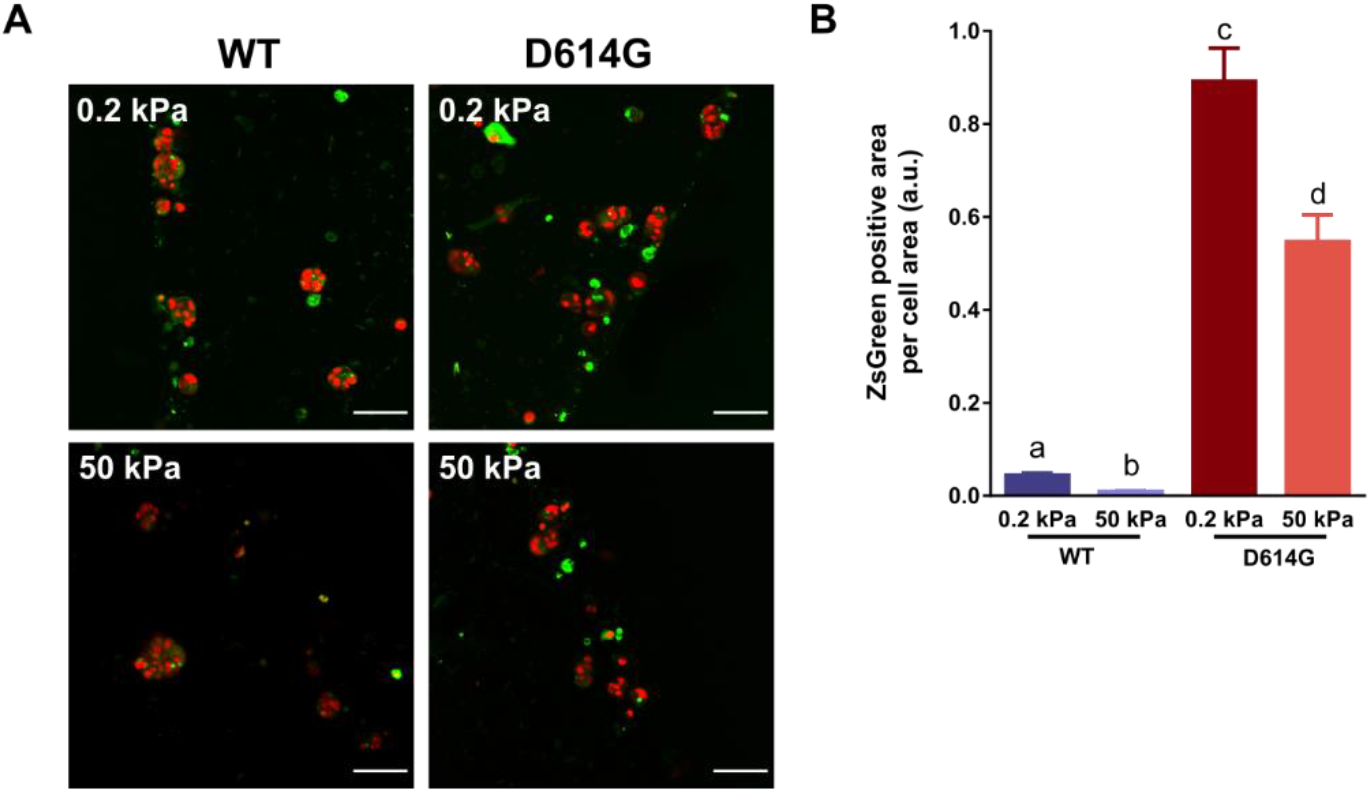
PV infection (WT and D614G) of Calu-3 lung cells on soft (0.2kPa) and stiff (50kPa) hydrogels. (A) Fluorescent images showing cells (red, DRAQ5) expressing ZsGreen (green) after successful infection. Scale bars = 100 μm. (B) Cells on softer substrates are preferably infected and increased infection rates are found with the D614G variant on both scaffolds. Different letters above the bar (a-d) denote statistically significant difference (p<0.05) (mean ± SEM, N=3/n=42 for 0.2kPa/WT, N=4/n=70 for 0.2kPa/D614G, N=3/n=49 for 50kPa/WT, N=4/n=67 for 50kPa/D614G).

### PV infections increase on sparse fibers

In addition to fibrotic stiffening of the lung parenchyma with age, increased ECM fiber density also occurs in stiffer (older) lungs [49], [50]. It has been further speculated that pathogens can better stick to the ECM and thus increase the infection probability [51]. However, the collagen hydrogel system that we used in Figure 2 does not de-couple stiffness from fiber density. Thus, we used electrospun fiber mats in addition, which are well known to mimic the geometrical properties of ECM fibers. Specifically, we used polycaprolactone (PCL) fiber matrices, which have been previously used as *in vitro* lung models [52], [53]. We used collagen-coated electrospun PCL fibers that were either sparse or dense by changing the collection time (**Figure 3A**) with all other parameters being identical. We found an average of 2.5x more fibers on the dense scaffolds (**Figure 3B**), while cells formed smaller cell clusters (**Figure 3C**). Cells on sparse fibers showed more infection events for the WT and D614G variants (**Figure 3D, E**). Interestingly, the D614G variant lost its increased efficiency relative to the WT on both sparse and dense electrospun mats, suggesting nanofiber scaffolds on glass have unique features.

**Figure 3.**
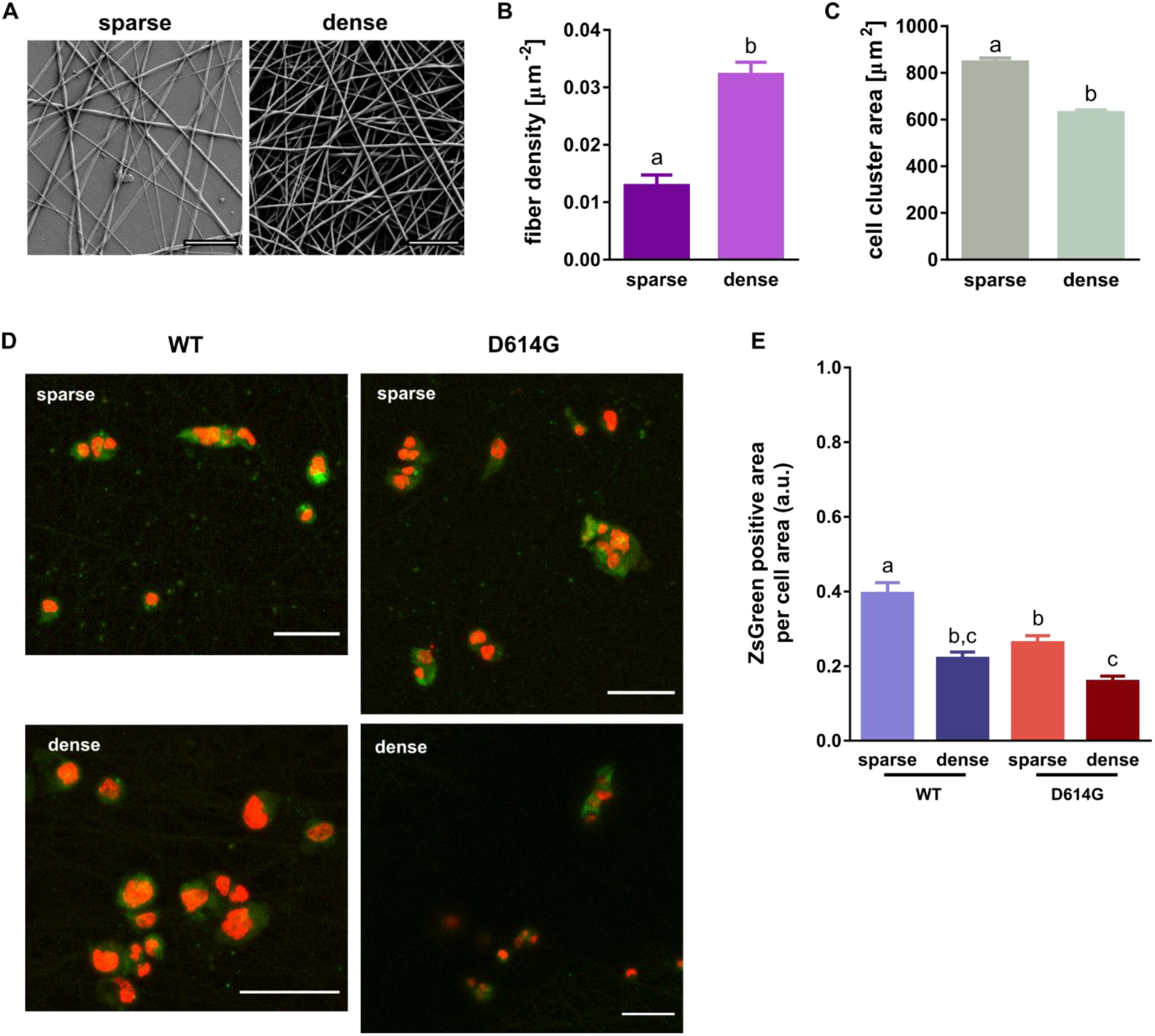
PV infection (WT and D614G) of Calu-3 lung cells on sparse and dense PCL fiber matrices. (A) SEM images showing the electrospun PCL fibers. Scale bars = 50 μm. (B) Fiber density is measured from the number of fibers per area. (C) Cells form clusters of different sizes on the two scaffolds. (D) Fluorescent images showing cells (red, DRAQ5) expressing ZsGreen (green) after successful infection. Scale bars = 100 μm. (E) Increased infection rates are found on sparse scaffolds, while the D614G variant is less infectious on the scaffolds. Different letters above the bar (a-c) denote statistically significant difference (p<0.05) (mean ± SEM, N=3/n=78 for sparse/WT, N=4/n=116 for sparse/D614G, N=4/n=125 for dense/WT, N=4/n=121 for dense/D614G).

### ACE-2 expression is modulated by cell-material interaction

To further evaluate the potential reasons underlying the stiffness-dependent infection of Calu-3 cells with SARS-CoV-2 PVs, a natural question is whether the amount of ACE-2 receptor expression varied among the cells on different substrates. Using immunofluorescence, we found a lower level of ACE-2 in Calu-3 than ACE-2-expressing HEK 293T on TCPS (**Figure 4A, B**), which is not surprising since the HEK 293T are genetically engineered to overexpress ACE-2. ACE-2 expression occurred in clusters in both cell types, albeit more so in the Calu-3. When we compared the ACE-2 expression on the hydrogel substrates, we observed a significant decrease in ACE-2 expression with increasing stiffness (**Figure 4C, D**). This correlates well with decreased expression of ACE-2 found in lungs of older rats [54], which has also been shown in other tissues in mice [55] and agrees with our finding that cells on softer substrates showed more PV infection. Additionally, Calu3 cells express more ACE-2 on dense scaffolds than sparse fibers (**Figure 4E, F**) in an opposing trend to cell cluster size (**Figure 3C**). However, the overall ACE-2 expression is closer to that on glass, which aligns with the elastic modulus of 3800 kPa for electrospun PCL [56], being more similar to glass.

**Figure 4.**
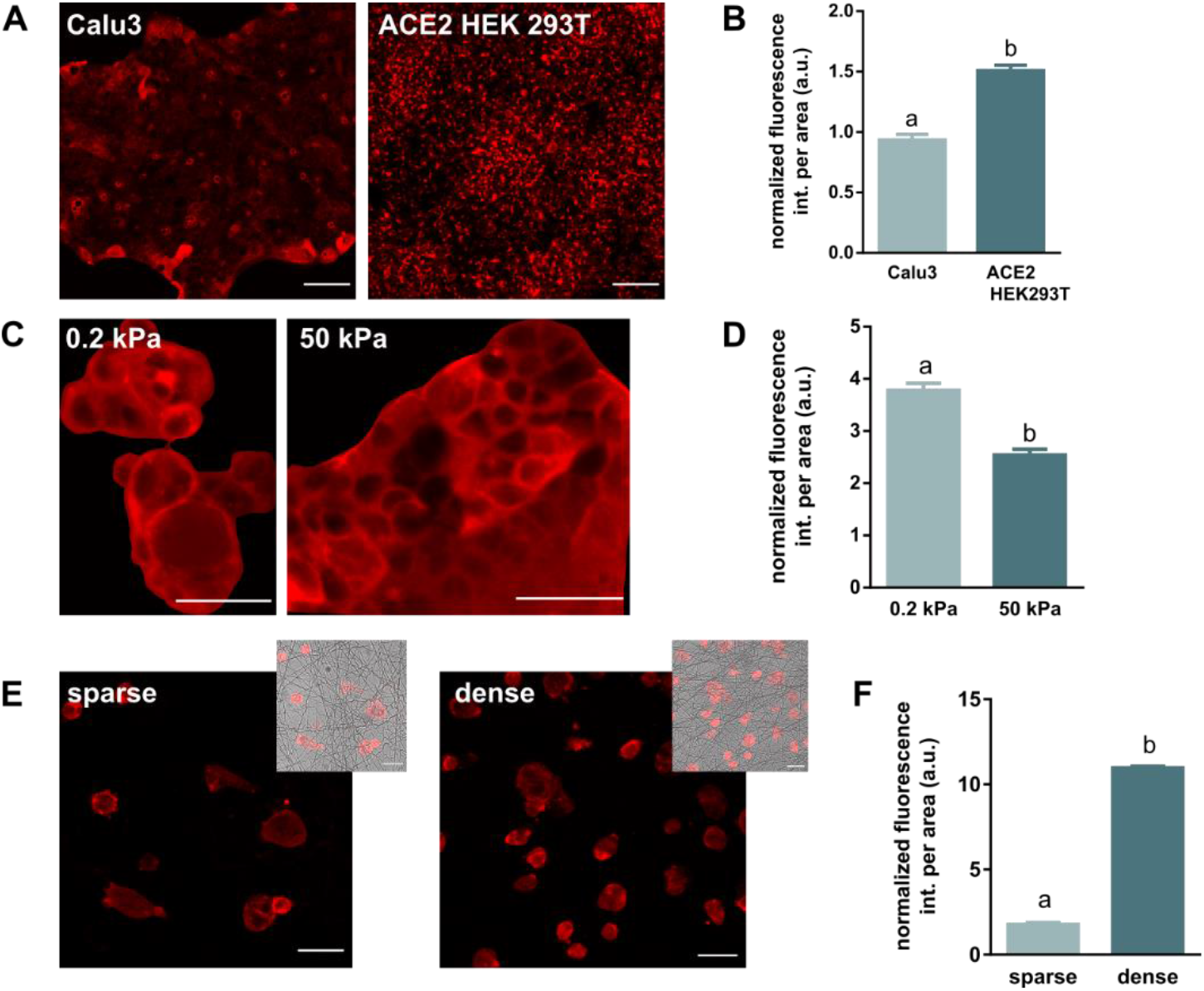
ACE-2 expression on different culture surfaces. (A) ACE-2 expression in Calu-3 lung cells and ACE-2 HEK 293T on TCPS. Fluorescent images show ACE-2 immunofluorescence (red, anti-ACE-2-Alexa546). Scale bars = 50 μm. (B) Fluorescent intensity per area normalized to Calu3 on TCPS. (C, D) Higher expression rates are found in Calu-3 on softer substrates (0.2 kPa) compared to stiffer surfaces (50 kPa). (E, F) Calu-3 on denser matrices express more ACE-2. Different letters above the bar (a-b) denote statistically significant difference (p<0.05) (N=1 for hydrogels/fibers, N=2 for TCS, n=32 for Calu-3 on TCPS, n=15 for ACE-2 HEK 293T, n=20 for 0.2kPa and 50kPa, n=19 for sparse, n=18 for dense).

Calu-3 lung cells grown on softer surfaces and incubated with PVs showed increased infection compared to stiffer hydrogels and TCPS by 4.8-fold and 1.6-fold, respectively (**Figure 1B**). A possible mechanism underlying this mechanosensitivity of viral infection is that cells can sense the local mechanical environment and respond by changing their mechanics or expression profiles. For example, cells have been shown to alter membrane stiffness [57], and receptor expression [58] led to modified uptake efficiency for antigens and nanoparticles [58]. Svenja *et al*. demonstrated that dendritic cells cultured on soft surfaces (2 kPa) expressed a high amount of surface lectin receptors (like mannose receptor and DC-SIGN) in comparison to the expression on stiff surfaces (50 kPa). Changes to the substrate stiffness significantly regulated the cells’ ability to recognize and internalize foreign sugar-based antigens [58]. On the other hand, Wang *et al*. illustrated that spread cells on stiffer surfaces internalized a significantly lower amount of nanoparticles per unit area than lesser spread cells on softer surfaces [59]. Similar behavior was observed for the interaction of SARS-CoV-2 with different macrophages, where the phenotype with the largest cell membrane stiffness showed the lowest viral uptake [47]. We note, ECM substrate stiffness has been shown to affect cell phenotype by inducing an epithelial-to-mesenchymal transition in certain cells [60], [61], which itself may also modify viral uptake. However, such an effect is not expected to play an important role here because Calu-3 do not transition to a mesenchymal phenotype, even under TGF-β1 stimulation [62].

Similar to soft or stiff hydrogels, ECM fiber density also significantly led to changes in ACE-2 expression, with dense fiber mats leading to increased expression. In this case, the level of ACE-2 expression did not correlate with infection levels as Calu-3 cells on sparse fibers always showed more infection and had lower ACE-2 expression. Cell spreading, which in turn affects membrane stiffness and endocytosis [21] (as we discuss below), is known to depend on fiber density for natural ECMs and electrospun scaffolds [63]. Analysis of cell cluster areas of Calu-3 cells on the matrices shows decreased spreading on dense fibers compared to fiber matrices (**Figure 3C, D**), presumably, resulting in higher membrane tension due to more spreading on sparse fibers. Using a biophysical model, Wang *et al*. showed initial adhesion of spherical viral particles (resembling SARS-CoV-2 and Zika) on host membranes depends on bending stiffness and membrane tension, where highly tensed membranes showed weak adhesion between viruses and cell membranes [64]. Huang *et al.* found that particle uptake per cell area was reduced for more spread cells, but that total uptake increased overall compared to less spread cells. This effect was modeled via energetic costs of binding, uptake, and cell spreading [21]. Taken together, this suggests a reason why larger, more spread clusters of cells (but not too spread out) take up more PVs compared to smaller clusters.

Our results show that the virus infection is more dependent on ACE-2 expression for the hydrogels, with cells on the 0.2kPa substrate expressing more ACE-2 and exhibiting higher initial infection than on the 50kPa substrate. On the other hand, on the fiber matrices, the cell surface area seems to become a more important factor, which might explain why the D614G variant loses its potency.

## Conclusion

The present work reports experimental evidence, from a controlled study, on the influence that local biomechanical properties of the tissue environment have on cellular uptake of SARS-CoV-2 viral particles. Successful SARS-CoV-2 infections seem to rely strongly on a combination of appropriate tissue elasticity, with softer tissues facilitating higher infection rates (especially true for the D614G variant) and sufficient ACE-2 expression. These findings indicate that worse clinical outcomes in older patients are not necessarily caused by existing biomechanical changes in lung tissue parenchyma that drive early infection events but possibly by other effects. Importantly, our work does not specifically address the post-infection replication of the virus and immune response, which could indeed be modulated by ECM microenvironment and tissue stiffness and should be studied further.

## Acknowledgments

We thank the Jesse Bloom lab for the kind provision of the SARS-CoV-2 pseudotyped lentivirus plasmids and associated cell lines. S.H.P. acknowledges support from the Human Frontier in Science Program (RGP0045/2018), the Welch Foundation (F-2008-20190330), and Texas 4000 Seed grant funding. A. P. was supported by the Swedish research council (2019-00682) and the Barbro Osher Endowment (2019-0124). K.N.D. received funding from the Welch Foundation (F-1390).

## Author Contribution

A.P., S.K., T.S.K., M.P., and A.L.B. performed the experiments and data analysis. A.P., S.K., and S.H.P. conceptualized, designed the study and wrote the manuscript. J.Z. and K.N.D. provided important discussion and materials. All authors contributed edits to the manuscript and approved of its final form.

