## Supplemental data and videos for "Biomechanical dependence of SARS-CoV-2 infections": SARS-CoV-2-biomechanics-SI.pdf

**Supporting Figures**

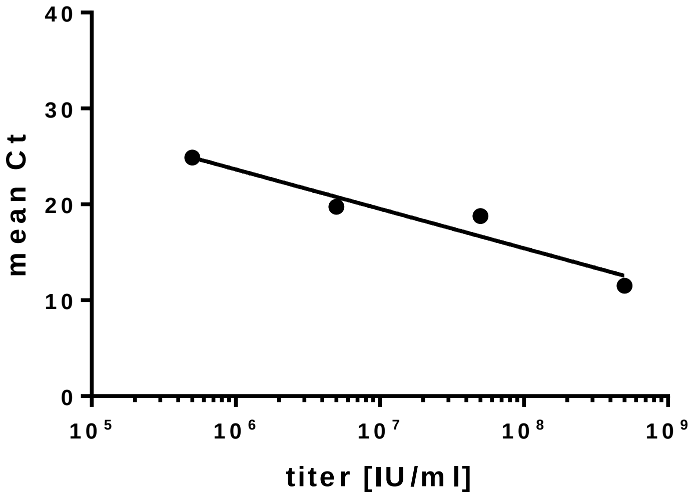


**Figure S1.** qPCR calibration of viral DNA content *versus* titer.

**Supporting Tables**

**Table S1.** Viral titer in different batches of pseudotyped lentivirus production as calculated from the standard calibration curve shown in Figure S3.

| S protein mutation | mean Ct | titer [IU/ml] |
|---|---|---|
| **WT** | 19.772 | 8.77E+06 |
| **D614G** | 22.199 | 2.25E+06 |

**Supporting Videos**

**Video S1.** ZsGreen expression in ACE2 HEK 293T infected with pseudotyped lentiviruses displaying VSV-G (viral envelope glycoprotein) during the first 72 hours (image spacing: 2 h). Scale bar 300 µm.

**Video S2.** ZsGreen expression in ACE2 HEK 293T infected with pseudotyped lentiviruses displaying WT S during the first 72 hours (image spacing: 2 h). Scale bar 300 µm.

**Video S3.** ZsGreen expression in HEK 293T infected with pseudotyped lentiviruses displaying VSV-G (viral envelope glycoprotein) during the first 72 hours (image spacing: 2 h). Scale bar 300 µm.

**Video S4.** ZsGreen expression in HEK 293T infected with pseudotyped lentiviruses displaying WT S during the first 72 hours (image spacing: 2 h). Scale bar 300 µm. No infection events can be observed.
